# Characterizing Mitochondrial Dysfunction Across Time in a Porcine model of Spinal Cord Injury

**DOI:** 10.64898/2026.02.05.704056

**Authors:** Olivia J Kalimon, Judee Grace E Nemeño, Candace L Floyd, Lonnie E Schneider

## Abstract

Spinal cord injury (SCI) can result in temporary or permanent alterations in sensory, motor, and autonomic functions as a result of primary mechanical damage to the spinal cord. Functional recovery is often limited due to persistent secondary injury mechanisms like inflammation, vascular breakdown, and cellular damage. Mitochondrial dysfunction is a key driver of secondary injury pathology, and while mitochondrial-targeted therapies have shown promise in rodent models of injury, functional improvements fail to translate to humans. Pigs are excellent models for understanding both the behavioral and molecular consequences of SCI because of their physiological similarity to humans, which could bridge the translational gap between rodent research and clinical implementation. To develop effective, mechanistic-based therapies, we must understand the molecular underpinnings of SCI using both male and female animal models with high translational fidelity at multiple time points after injury. To date, research on mitochondrial dysfunction following SCI has been limited to female rodent models measured acutely (6h-7d) after injury. Here, we studied mitochondrial dysfunction at three different time points in male pigs to establish a relative time course of mitochondrial impairment following SCI that may be therapeutically targeted to treat secondary complications of injury. We measured mitochondrial bioenergetic function and electron transport chain (ETC) complex activities, as well as qualified mitochondrial dynamics and oxidative damage acutely (2h), sub-acutely (24h), and chronically (9wk) after SCI in adult male pigs. The results show distinct patterns of mitochondrial dysfunction between time points with functional deficits occurring 2h post-SCI, increased mitochondrial fragmentation at 24h post-SCI, and mitochondrial recovery by 9wks post-SCI. These studies offer insight into mitochondrial changes across time in a clinically relevant animal model of SCI in hopes of bridging the translational research gap.

## Introduction

Spinal cord injury (SCI) is a debilitating disorder that results in loss of sensorimotor function following mechanical damage to the spinal cord. Individuals living with SCI can develop a wide array of secondary complications, like autonomic dysreflexia, spasticity, or neuropathic pain, that can make daily life very difficult [1-4] Primary mechanical injury to the spinal cord sets off a host of secondary injury mechanisms, such as excitotoxicity, inflammation, and mitochondrial dysfunction, that lead to cell death and scar formation which contributes to long-term complications and limits functional recovery following SCI [5]. The current treatment options for SCI include surgical intervention, physical therapy, hypobaric oxygen therapy, and pharmaceuticals [6, 7]. However, these treatment options do not “cure” SCI; these are only strategies to mitigate secondary complications of injury. To create effective therapies to improve functional recovery and reduce the prevalence and persistence of secondary complications after SCI, researchers must understand the underlying mechanisms following SCI across time using more translational animal models.

Mitochondria have been dubbed the “powerhouses of the cell” due to their crucial roles in energy production, calcium homeostasis, and oxidative stress, each of which serve to maintain cell viability. Mitochondrial dysfunction is a major cause and consequence of secondary injury following damage to the central nervous system and, as such, mitochondria have been targets of therapeutic interest [8-10]. Following primary mechanical damage to the spinal cord, there is increased excitotoxic glutamate released that drives an influx of calcium into the cell [11, 12]. Mitochondrial calcium influx depletes mitochondrial membrane potential and, if it cannot be restored due to excessive intracellular calcium resulting from injury, mitochondria can undergo permeability transition and release pro-apoptotic/necrotic factors to trigger cell death [13-15]. Reactive oxygen/nitrogen species (ROS/RNS) can also impair mitochondrial respiratory functions, that can further exacerbate oxidative stress and worsen secondary injury following SCI [16, 17]. Studies within the fields of both traumatic brain injury and SCI have found that mitochondrial targeted therapies can improve mitochondrial bioenergetics and limit oxidative damage to reduce gross tissue damage and improve functional recovery [8, 9, 18-22]. Researchers have even explored the efficacy of mitochondrial transplantation in promoting functional recovery following SCI [23-25]. Further, studies measuring mitochondrial dysfunction across time have shown that mitochondrial impairment does not always occur in a linear fashion following injury [17, 26-28]. This suggests that we need a detailed understanding of secondary injury and recovery across time so that we may best treat the people living with these conditions.

Most of what is known about mitochondrial impairment after SCI has been performed in rodent models of injury; and more specifically, female rodent models of injury [17, 21, 22, 29]. However, approximately 78% of new SCI cases occur in men, which suggests that further research should be done to elucidate the post-SCI mitochondrial mechanisms in male animal models. To create effective, mechanistic-based therapeutics, we must use animal models with increased translational fidelity that expands beyond one sex and one timepoint after injury. This present study was designed to characterize mitochondrial dysfunction in castrated male pigs at the acute (2h), subacute (24h), and chronic (9wks) times following contusion-compression SCI. Compared to rodents, pigs are more similar to humans with respect to their size, neural organization, blood flow, metabolism, and immune response which makes them excellent translational models to study the molecular and behavioral consequences of SCI. We measured mitochondrial bioenergetics and electron transport chain (ETC) complex activities, as well as the expression of proteins related to mitochondrial dynamics (i.e. fission/fusion and mitophagy/biogenesis) and oxidative damage.

## Materials and Methods

### Animals and Experimental Design

Adult castrated and gonad intact male Yucatan pigs were randomly assigned to naïve control, laminectomy control (LAM) or contusion-compression spinal cord injury (SCI) groups. For acute (2h) groups, pigs remained anesthetized until 2h post-surgery until humane euthanasia. For subacute (24h) groups, pigs were allowed to recover from anesthesia in an acute care setting until humane euthanasia at 24h post-surgery. For both acute and subacute timepoints, one LAM and one SCI were generated on each surgery day, and group numbers were accumulated across multiple days. For chronic (8-10wk) SCI group, pigs were allowed to recover for 7-10 days in an acute care setting before returning to modified group housing. To limit the use of animals, there was no chronic LAM control group; however, samples were collected from SCI “naïve” pigs (i.e. cerebrospinal fluid port) as uninjured controls for analysis at this time point. For the chronic time point, two SCI surgeries were performed on each surgery day and group numbers were accumulated across multiple days. Chronic SCI pigs underwent weekly behavioral testing for another study (*data not shown*) before humane euthanasia at 8-10 weeks post-SCI. All in vivo animal procedures including handling and acclimation, surgery, behavioral training and testing, daily post-operative care, humane euthanasia, tissue collection, and tissue processing were performed at Emory University.

### Contusion-Compression Spinal Cord Injury and Associated Cystotomy Surgical Procedures

All experiments were approved by and conducted in accordance with the Emory University Institutional Animal Care and Use Committee (IACUC), the Office of Laboratory Animal Welfare Guide for the Care and Use of Laboratory Animals, the U.S. Government Principles for the Utilization and Care of Vertebrate Animals Used in Testing, Research and Training, and the Animal Welfare Act. Pigs were fasted for 12h prior to surgery. Immediately prior to surgery, pigs were anesthetized with an intramuscular injection of Telazol (4.4 mg/kg), ketamine (2.2 mg/kg) and xylazine (2.2 mg/kg) and were then intubated with an endotracheal tube to be maintained on 1-4% isoflurane anesthesia. All surgical procedures were performed in a designated large animal operating suite under strict aseptic conditions with continuous monitoring of heart rate, respiratory rate, end tidal carbon dioxide and oxygen saturation. Lidocaine was administered to the perioperative area by subcutaneous injection prior to incision. Temperature was maintained (∼38-39°C) during surgical procedures. Hydration was maintained with IV saline during surgical procedures.

For the induction of SCI, overlaying skin and muscle layers were dissected from the spinal column. A laminectomy was performed at the 10^th^ thoracic vertebrae (T10) and the periosteum was removed to expose the spinal cord. The weight drop apparatus was affixed to the spinal column using multiaxial pedicle screws (3.5×2.4mm screw, Medtronic). 3.2mm titanium rods were affixed to the screws to secure the weight drop guidance system to the spinal column. A short acting paralytic (1mg/kg succinylcholine) was administered intravenously to inhibit motor discharge upon impact to the spinal cord and one breath was held upon dropping the impactor (50g) onto the spinal cord. The impactor was dropped from 20cm height, which corresponds to a displacement of ∼2mm and a force of ∼35N. After one minute, an additional 100g weight was placed onto the impactor to model spinal cord compression for 5 minutes. After the 5m, the weights and guidance system were removed and the holes within the pedicles were sealed with sterile bone wax. Injury was confirmed with visualization of bruising on the spinal cord without disruption of the dura. The LAM group received the same procedures, without impact and subsequent compression of the cord. Upon completion of the procedure, the incision was closed in layers.

All pigs that underwent spinal surgery (i.e. LAM/SCI groups) received a cystotomy and implantation of a urinary catheter. This procedure is performed to ensure bladder voiding during the transient state of spinal shock following SCI, where pigs experience bladder dysfunction and cannot urinate on their own. Male pigs have a sigmoid flexure with an acute bend in the distal loop in the body of the penis that does not allow for clean intermittent catheterization, necessitating the placement of an indwelling catheter in the bladder. A midline posterior abdominal incision was made, and the bladder was subsequently exposed. A purse-string suture was made within the bladder wall, and a stab incision was made in the center of the purse-string. A foley catheter (14-18FR) was immediately placed in the incision, the balloon was inflated, and the purse-strings were secured. The urinary catheter was secured to a sterile collection bag that was emptied when necessary, using aseptic technique. The abdominal incision was closed in layers.

Upon completion of all surgical procedures, after all incisions were closed, anesthesia was discontinued, and the pigs were extubated once spontaneous respiration occurred. The acute (2h) groups were maintained on anesthesia until humane euthanasia at 2h post-surgery. The subacute (24h) and chronic groups (9wks) groups were recovered in a warmed room and were then moved to the specialized acute care setting once awake and alert. The subacute groups remained in the acute care setting until humane euthanasia at 24h post-surgery. The chronic group remained in the acute care setting for ∼10d post-SCI until returning to modified, standard housing. In the acute care setting, pigs were individually housed in the lower half of an XL clamshell dog kennel containing memory foam dog bed, blankets, towels, pillows, and enrichment items, which were changed daily. Pigs were handfed and watered with frequent health checks and repositioned to minimize the prevention of pressure sore formation. Feces were manually removed, and the skin was kept clean and dry by trained staff. While in the acute care setting, pigs receive visual, olfactory, and auditory stimuli from other pigs. Once robust voluntary micturition via the urethra was observed (∼7-10d post-injury), the urinary catheter was removed, and the pigs returned to modified, standard group housing. Modifications include lowered food, water, and enrichment items.

### Tissue Dissection and Mitochondrial Preparation

At the specified endpoints, the pigs were sedated, anesthetized, and then humanely euthanized by barbiturate overdose, according to the Veterinary Medical Association guidelines for euthanasia of animals. For the 2h time point, one LAM Control and one SCI pig were euthanized each day of experimentation (total 3 days of experimentation). For the 24h time point, one Naïve Control (naïve to all surgical procedures) and one SCI pig were euthanized each day of experimentation (total 3 days of experimentation). For the 9wk time point, one uninjured control (naïve to spinal surgery [represented as open circle in all figures]) and three SCI pigs were euthanized on 1 day of experimentation. Pigs underwent cardiac perfusion with ice-cold phosphate buffered saline (0.1M PBS) using a low-pressure pump (Model 170DM5, Stenner Pump Company) set to 200 mL/min. The spinal cord at T9-10 was visualized and then a ∼1cm region was excised. For SCI groups, the 1cm region of spinal cord was centered on the SCI epicenter (T10) and the surrounding penumbra. After dissection, the dura mater was carefully removed and then the spinal cord was homogenized in 6mL mitochondrial isolation buffer (215mM mannitol, 75mM sucrose, 0.1% BSA [w/v], 1 mM, and 20 mM HEPES pH 7.2 with KOH) using a glass dounce tissue homogenizer. One aliquot (2 mL) of tissue homogenate was reserved and stored at -80°C for oxidative stress protein analyses while the remaining samples (4 mL) were processed for mitochondrial analyses. Crude mitochondrial pellets were isolated using the differential centrifugation method, described as follows. Homogenates were centrifuged twice at 1,300xg for 3 m at 4°C, reserving the supernatants after the first spin and then again after resuspending the pellets and centrifuging for the second time. The reserved supernatants were centrifuged at 13,000xg for 10 m at 4°C. The pellets were resuspended in isolation buffer to ∼500 µL volume and then subjected to nitrogen disruption at 1,200 psi for 10m on ice. The samples were transferred to a clean 1.5 mL Eppendorf tube and centrifuged at 13,000xg for 10m at 4°C to achieve the crude mitochondrial pellet. Protein concentration was estimated using the Pierce BCA Protein Assay Kit (Thermo Fisher, Cat. 23225).

### Bioenergetics

Mitochondrial bioenergetics were measured in the presence of substrates, inhibitors, and uncouplers of the mitochondrial electron transport chain (ETC). Oxygen consumption rate (OCR) was measured using the Seahorse XFe24 Analyzer (Agilent). Isolated mitochondria (60µg) were diluted in respiration buffer (125 mM KCl, 2 mM MgCl_2_, 2.5 mM KH_2_PO_4_, 20 mM HEPES, 0.1% BSA [w/v], pH 7.2 with KOH) before being loaded onto the Seahorse XFe24 cell culture microplates (Agilent). Plates were centrifuged at 2204 x g for 10m at 4°C to ensure mitochondria remained at the bottom of the well. Warmed (37°C) respiration buffer was loaded slowly onto the top of each well immediately prior to running the assay. Stocks for mitochondrial substrates, inhibitors, and uncouplers were diluted in respiration buffer (without BSA) and then loaded into the ports to assess the different mitochondrial respiration states. State III respiration, or adenosine triphosphate (ATP) production-linked respiration, was measured in the presence of pyruvate (5 mM), malate (2.5 mM), and adenosine diphosphate (ADP; 2 mM). State IV respiration, or proton leak respiration, was measured in the presence of the ATP synthase inhibitor, oligomycin (2.5 µM). State V_CI_, or uncoupled respiration driven primarily by ETC Complex I, was measured in the presence of the mitochondrial uncoupler, carbonyl cyanide m-chlorophenylhydrazone (CCCP; 4µM). State V_CII_, or uncoupled respiration driven by ETC Complex II, was measured in the presence of the Complex I inhibitor, rotenone (0.1 µM), and the Complex II-specific substrate, succinate (10 mM). For the 2h and 24h time points, one control and one SCI pig were run on one Seahorse plate per day, with 3d of experimentation per time point (n=3/group). For the 9wk time point, one control and three SCI pigs were run on one Seahorse plate.

### ETC Complex Activities

Mitochondrial electron transport chain (ETC) complex activities were measured in Seahorse XFe24 Analyzer (Agilent) in the presence of substrates and inhibitors of the ETC in isolated mitochondria that had been previously frozen, as described previously with some modifications for use with the Seahorse XFe24 [30]. Isolated mitochondria (30 µg) were diluted in respiration buffer before being loaded onto the Seahorse XFe24 cell culture microplates (Agilent). Plates were centrifuged at 2204 x g for 10m at 4°C to ensure mitochondria remained at the bottom of the well. Warmed (37°C) respiration buffer (with BSA) containing 10 µM cytochrome C, 30 µM nicotinamide adenine dinucleotide (NADH), and 10 µg/mL alamethicin was loaded slowly onto the top of each well immediately prior to running the assay. Stocks for mitochondrial substrates and inhibitors were diluted in respiration buffer (without BSA) and then loaded into the ports as follows: (A) 0.8 µM rotenone and 10 mM succinate, (B) 1 µM antimycin A, (C) 20mM ascorbate and 5mM N,N,N’,N’-tetramethyl-p-phenylenediamine (TMPD), (D) 549.3 mM sodium azide. 2-3 oxygen consumption rate (OCR) readings were taken to ensure the full activity was measured after the addition of NADH, as well as after each port addition. Complex I activity was calculated by subtracting the antimycin A reading from the NADH reading. Complex II activity was calculated by subtracting the antimycin A reading from the rotenone/succinate reading. Complex IV activity was calculated by subtracting the azide reading from the ascorbate/TMPD reading.

### Mitochondrial Dynamics Western Blot

Mitochondrial dynamics (i.e. fission, fusion, mitophagy, biogenesis) were assessed by Western blot. Mitochondrial samples were loaded onto a 4-20% Criterion TGX gel (BioRad, Cat. 5671095) and subjected to gel electrophoresis at 120V. Proteins were then transferred to a PVDF membrane (BioRad Midi format Trans-Blot Turbo Transfer Pack, Cat. 1704157) using the BioRad Trans-Blot Turbo Transfer System according to the preprogrammed Mixed MW setting (2.5A, up to 25V). Membranes were then prepared for Revert Total Protein stain, described previously with some modifications [31]. Membranes were dried at 37°C for 30m, then rehydrated in 99.8% methanol for 1m, followed by incubation in PBS for 5m. Membranes were incubated in diluted (1:4) Revert 700 Total Protein Stain (Licor, 926-11011) for 10 minutes at room temperature and then washed 3×1m with Revert Wash Buffer. Membranes were imaged using the Odyssey CLx Imager, and then incubated in Revert Destaining Buffer for 7m shaking on a rocker. The membranes were then incubated in 5% milk in TBST Wash Buffer (1X tris buffered saline, 0.1% Tween 20) for 1h at room temperature, followed by overnight incubation at 4°C in primary antibody (Drp1 1:1,000 [abcam, ab184247], LC3B 1:2,000 [abcam, ab192890], OPA1 1:1,000 [Novus Biologicals, NB110-55290], OxPhos Human WB Antibody Cocktail 1:500 [Thermo Fisher, 45-8199], PGC1α 1:1,000 [abcam, ab191838]). The next day, membranes were washed 3×5m in TBST and then incubated in fluorescent secondary antibody (1:15,000 IRDye 680RD Goat anti-Mouse IgG [Licor, 926-68070] or 1:20,000 IRDye 800CW Goat anti-Rabbit IgG [Licor, 926-32211]) in 3% BSA for 1h at room temperature. Membranes were washed 3×5m in TBST and then imaged on the Odyssey CLx Imager. The sum signal intensity of each sample was quantified in Image Studio 6.0 (Licor) and then normalized to the sum signal intensity of total protein.

### Oxidative Stress Enzyme-Linked Immunosorbent Assay

Detection and quantification of malondialdehyde (MDA), 3-nitrotyrosine (3-NT), and protein carbonyl (PC) in spinal cord tissue homogenates were performed through enzyme-linked immunosorbent assay (ELISA) [32] according to the manufacturer’s instructions (MyBioSource; MDA ELISA kit: Cat. MBS742540, 3-NT ELISA kit: Cat. MBS754460, and PC ELISA kit: Cat. MBS2600294). Briefly, reserved tissue homogenates were sonicated at 40% amplitude at 20 pulse for 10 seconds. Protein concentration was estimated using the Pierce BCA Protein Assay Kit and all samples were made to the concentration. All plates and reagents were brought to room temperatures prior to starting the assay. Each standard concentration and each sample were analyzed in duplicate. Appropriate volumes of balance solution were added to each well loaded with homogenates only. Subsequently, 50 µL of antibody conjugates were dispensed to all wells then the plate was covered with an adhesive film and was incubated at 37°C for 1-1.5 hours. Thereafter, the plates were manually washed five times with their corresponding assay wash buffers. After each washing, the plates were inverted and gently blotted into absorbent papers to remove residual wash buffer. The Enzyme Conjugate and Color Detection Reagent were subsequently added at 50 µL each to all the wells then plates were covered and incubated at 37°C for 15-20 minutes. Stop Solution was added at 50 µL to each well including blank control and mixed thoroughly. The optical density (OD) values were determined at 450 nm using the Synergy H1 microplate reader (BioTek) and then concentration of protein (ng/mL) was calculated using the linear standard curve.

### Statistical Analysis

All statistical analyses were performed using GraphPad Prism 10.6.0. For bioenergetics experiments, one LAM and one SCI pig were run on one Seahorse plate each day of experimentation, and then data from all pigs (n=3/group) were pooled for analysis. Bioenergetic data for the 24h and 9wk Naïve groups were pooled together and then all groups were analyzed by one-way ANOVA with Dunnett’s multiple comparisons, where applicable (vs Naïve Control). For ETC complex activities experiments, reserved mitochondria (n=3/group) were run on one Seahorse plate per time point. Specifically, for the 9wk time point, the control group was composed of 1 spinal surgery Naïve Control (open circle) and 2 additional LAM Control animals (closed circles) (total n=3 control) to compare to 9wk SCI (n=3) on one Seahorse plate. ETC data for each time point was analyzed independently by one-tailed, unpaired t test. Western blot and ELISA experiments were designed for one-way analysis of variance (ANOVA). For Western blot experiments, all samples were run on the same gel per primary antibody. For oxidative stress ELISAs, all samples were run with the same standard curve per antibody plate. The Brown-Forsythe and Bartlett’s tests were employed to ensure homogeneity of variance of data. The Shapiro-Wilk test was employed to assess the normality of data. Data that did not pass normality assumptions were analyzed by the Kruskal-Wallis nonparametric test with Dunn’s multiple comparisons (vs Naïve Control), where appropriate. Data that passed the normality assumptions were analyzed by one-way ANOVA with Dunnett’s or Sidak’s multiple comparisons (vs Naïve Control or LAM Control), where appropriate. The specific post hoc analyses are listed in the figure legends.

## Results

### Bioenergetics

Mitochondrial respiration is crucial for maintaining all mitochondrial functions and is known to be impaired after injury. Previous studies in female rodents have found that mitochondrial respiratory dysfunction occurs as early as 12h post-SCI and persists at 3d post-SCI [17, 21, 22, 29]. To date, there have been no studies measuring mitochondrial respiratory function in male models of SCI. Here, we measured mitochondrial bioenergetics at the spinal cord injury epicenter of male pigs acutely (2h), sub-acutely (24h), and chronically (9wks) following contusion/compression SCI. Beginning 2h post-SCI, mitochondrial respiration is reduced across State III (Figure 1A), State V(CI) (Figure 1C), and State V(CII) ((Figure 1D) compared to Uninjured Control. Contrary to existing literature, mitochondrial respiration was not significantly different from Uninjured Control 24h post-SCI across any respiration states (Figure 1). Interestingly, State III respiration was increased 9wks post-SCI relative to Uninjured Control (Figure 1A).

**Figure 1.**
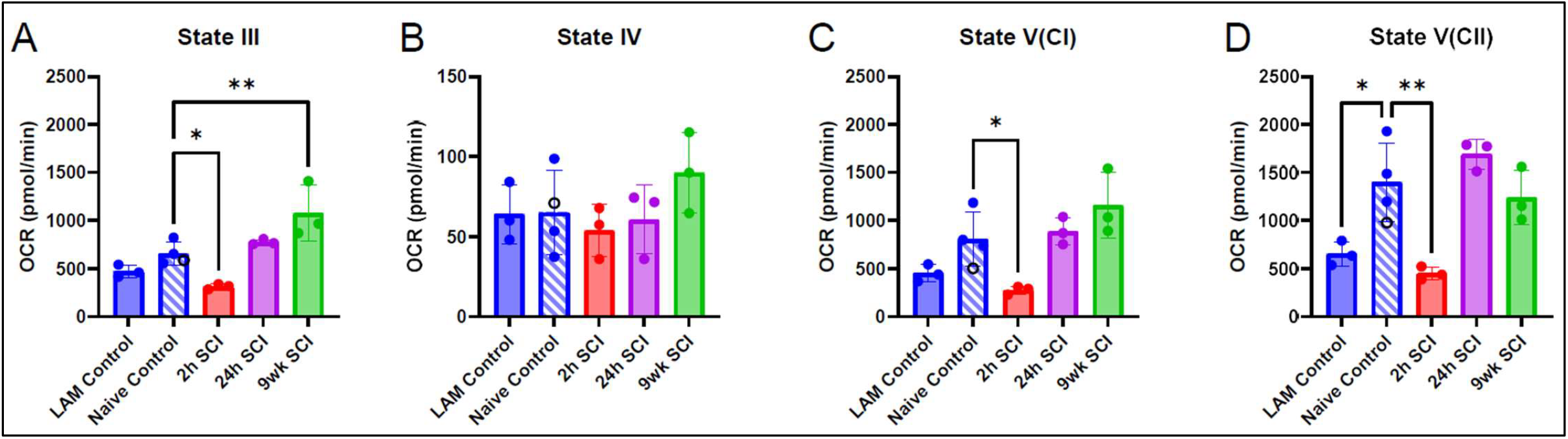
Mitochondrial bioenergetics is impaired 2h following SCI and appears to recover by 24h. (A) State III respiration was measured in the presence of pyruvate, malate, and ADP. (B) State IV respiration was measured in the presence of oligomycin. (C) State V(CI) respiration was measured in the presence of CCCP. (D) State V(CII) respiration was measured in the presence of rotenone and succinate. Data are represented as mean OCR ±SD and were analyzed by one-way ANOVA with Dunnett’s (A, C, D) multiple comparisons (vs Naïve Control). *p<0.05; **p<0.01.

### ETC Complex Activities

Mitochondrial bioenergetics represents how efficiently mitochondria function with membranes intact to give hints at the coupling or uncoupling of the oxidative phosphorylation (OXPHOS) process. To specifically examine Electron Transport Chain (ETC) complex activities, we performed another assay in mitochondrial samples that had been previously frozen (i.e. membranes are not intact). Conversely to bioenergetic data, Complex I, II, and IV activities 2h post-SCI were not significantly different from Uninjured Control (Figure 2A). At 24h post-SCI, each of the complexes had increased activity compared to Uninjured Control (Figure 2B). We also measured ETC complex activities in mitochondrial samples collected 9wk post-SCI but found no differences between SCI and Uninjured Control across any complexes (Figure 2C).

**Figure 2.**
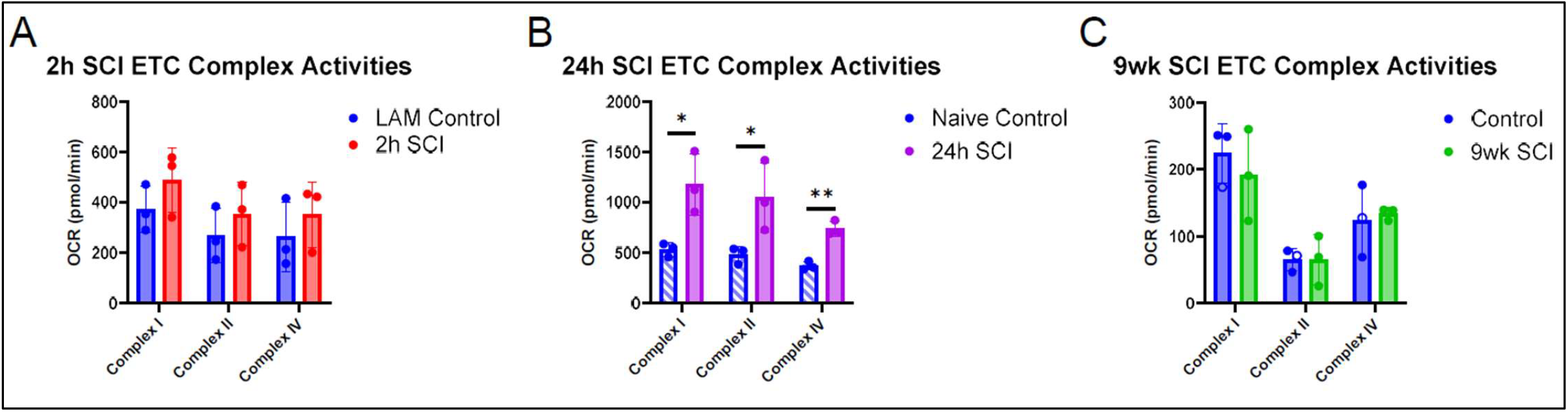
ETC complex activities are increased 24h post-SCI, while ETC complex activities are similar to control 2h and 9wk post-SCI. Complex activities were assessed independently by time point: (A) 2h post-SCI, (B) 24h post-SCI, and (C) 9wks post-SCI. Data are represented as mean OCR ± SD. Each complex for each time point were analyzed independently by one-tailed unpaired t test. *p<0.05; **p<0.01.

### Mitochondrial Dynamics

Balanced mitochondrial dynamics are essential for maintaining both mitochondrial and overall cellular health. When mitochondria are damaged, they undergo fission to separate the damaged mitochondrion from the healthy mitochondrial network. That damaged mitochondrion then undergoes mitophagy and is recycled to create a new mitochondrion through mitochondrial biogenesis. The new, healthy mitochondrion is then fused with the mitochondrial network to resume function. This mitochondrial quality control mechanism occurs normally throughout the body, even in the absence of injury, but has been shown to be upregulated after traumatic CNS injuries. In these present studies, we found expression of the mitochondrial fission protein, Drp1, to be significantly increased at 2h post-SCI and significantly decreased at 9wks post-SCI compared to Uninjured Control (Figure 3B). Expression of the mitochondrial fusion protein, OPA1, was significantly decreased at 24h post-SCI relative to Uninjured Control (Figure 3C). PGC1α, the transcriptional regulator of mitochondrial biogenesis, showed increased expression at 9wks post-SCI relative to Uninjured Control (Figure 3D). LC3B is a crucial protein for mitophagy with two distinct forms: LC3B-I (cytosolic, inactive form) and LC3B-II (lipidated, active form). While there were no differences in LC3B-II expression between the groups (Figure 3F), there was decreased expression of LC3B-I 9wks post-SCI relative to Uninjured Control (Figure 3E).

**Figure 3.**
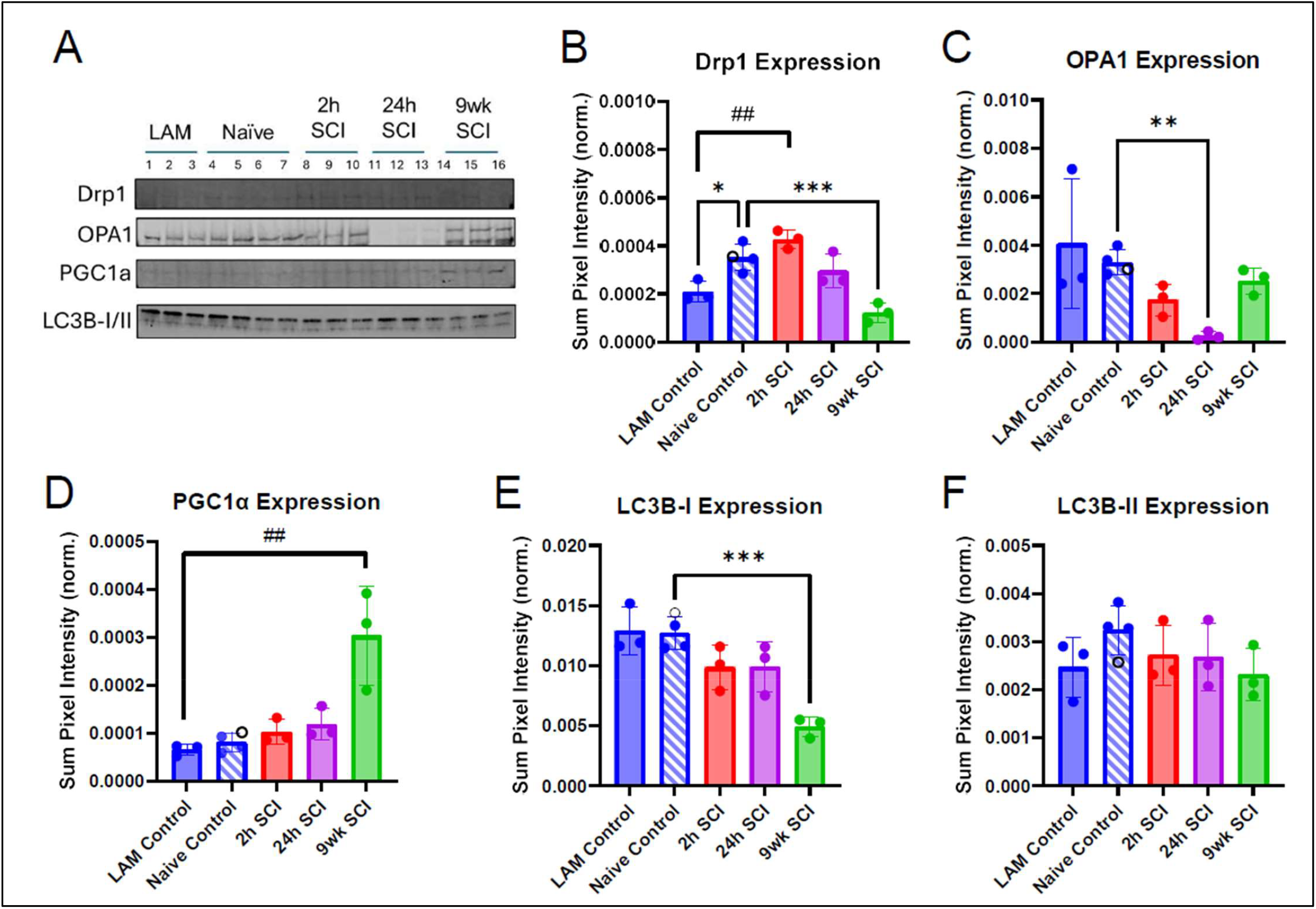
Mitochondrial turnover occurs acutely post-SCI, but mitochondrial recovery occurs chronically post-SCI. (A) Representative images of mitochondrial dynamics Western blot bands from isolated mitochondrial samples. (B) Expression of the mitochondrial fission protein Drp1. (C) Expression of mitochondrial fusion protein OPA1. (D) Expression of the transcriptional regulator of mitochondrial biogenesis, PGC1α. (E-D) Expression of the cytosolic and membrane-bound mitophagy protein LC3B-I and LC3B-II, respectively. Data are represented as mean ±SD sum signal intensity normalized to total protein. Data that passed normality assumptions were analyzed by one-way ANOVA (B, E, F) with Sidak’s multiple comparisons, where appropriate. Data that did not pass normality assumptions were analyzed by Kruskal-Wallis test (C, D) with Dunn’s multiple comparisons, where appropriate. Vs Naïve Control *p<0.05; **p<0.01; ***p<0.001. Vs LAM Control ^##^p<0.01.

In these studies, we also measured protein expression of proteins involved in OXPHOS (i.e. ETC Complex I, ETC Complex IV, and ATP Synthase, or Complex V). We found increases in Complex I (Figure 4B) and Complex IV (Figure 4C) expression, while Complex V (Figure 4D) expression was decreased 24h post-SCI relative to Uninjured Control. There were no other significant differences in OXPHOS protein expression at other time points.

**Figure 4.**
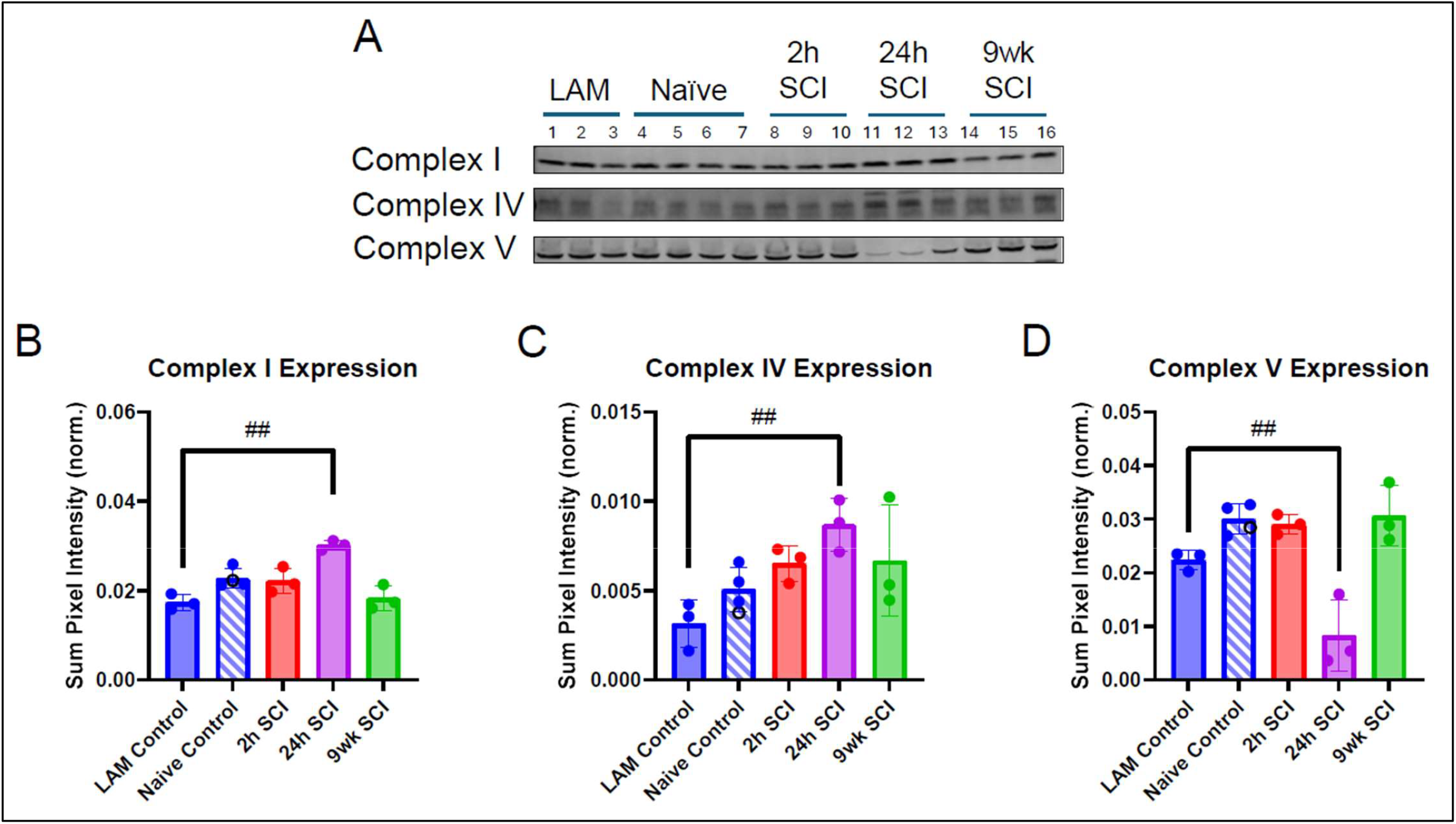
Protein expression of mitochondrial OXPHOS complexes is most variable 24h post-SCI. (A) Representative images of OXPHOS Western blot bands from isolated mitochondrial samples. (B) Expression of ETC Complex I. (C) Expression of ETC Complex IV. (D) Expression of Complex V (i.e. ATP Synthase). Data are represented as mean ±SD sum signal intensity normalized to total protein. Data that passed normality assumptions were analyzed by one-way ANOVA (C, D) with Dunnett’s multiple comparisons. Data that did not pass normality assumptions were analyzed by Kruskal-Wallis test (B) with Dunn’s multiple comparisons. Vs LAM Control ^##^p<0.01.

### Oxidative Damage

One well-known consequence of CNS injury is oxidative stress. Mitochondria are both generators and targets of reactive oxygen/nitrogen species (ROS/RNS) and, if not quenched by local antioxidants, these free radicals can bind to DNA, proteins, or lipid membranes to exacerbate damage following injury. The protein marker of RNS damage, 3-nitrotyrosine (3-NT), was increased beginning 2h post-SCI, and persisted through 9wks post-SCI relative to Uninjured Control (Figure 5A). There were no significant differences in relative quantity of protein carbonyl (PC), the marker of ROS damage (Figure 5B). Malondialdehyde (MDA) expression, a marker of lipid peroxidation, was significantly reduced 24h post-SCI relative to Uninjured Control (Figure 5C).

**Figure 5.**
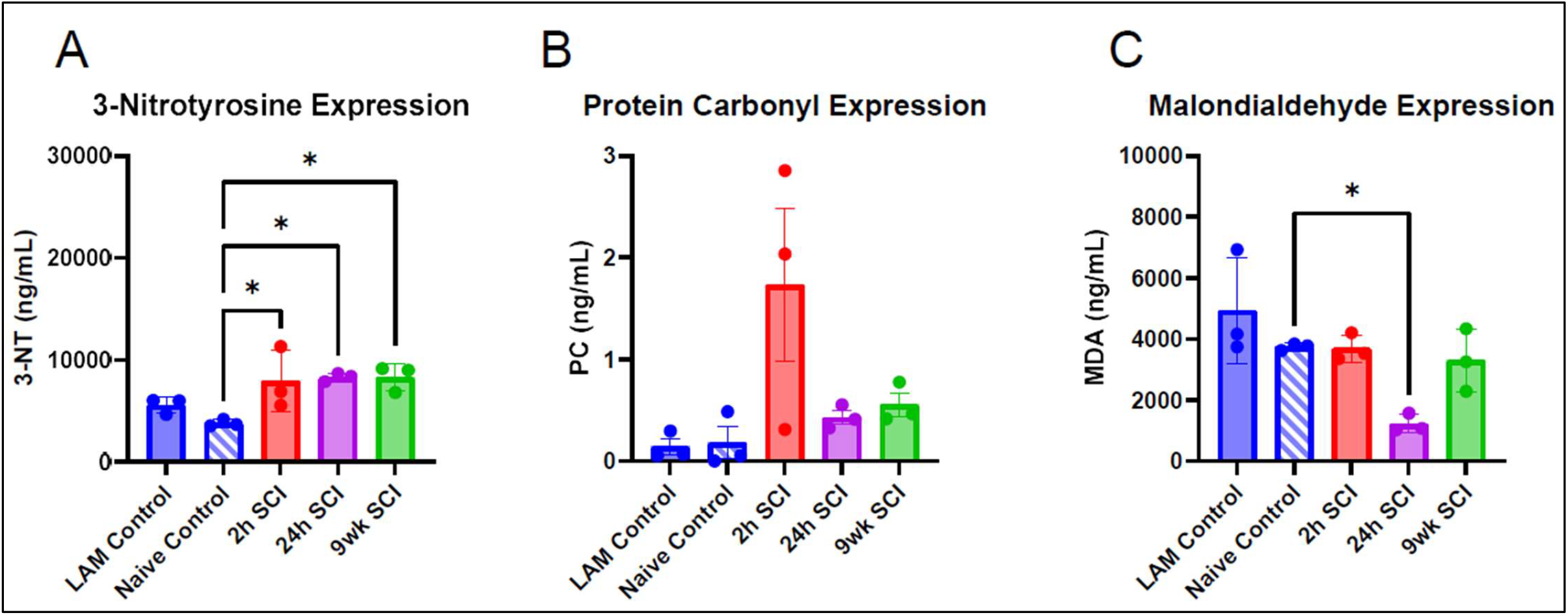
There is significant reactive nitrogen species damage in the spinal cord 2h post-injury that persists at 9wks post-SCI. Protein quantities within spinal cord homogenates were measured by ELISA. (A) Quantity of the reactive nitrogen species damage marker, 3-nitrotyrosine (3-NT). (B) Quantity of the reactive oxygen species damage marker, protein carbonyl (PC). (C) Quantity of the lipid peroxidation marker, malondialdehyde (MDA). Data are represented as mean (ng/mL) ± SD. Data that passed normality assumptions were analyzed by one-way ANOVA with Dunnett’s multiple comparisons (A, C). Data that did not pass normality assumptions were analyzed by Kruskal-Wallis test (B). Vs Naïve Control *p<0.05.

## Discussion /Conclusion

Healthy mitochondria are crucial for maintaining cell viability. Importantly, mitochondria have quality control mechanisms to maintain cellular health, even after injury. Mitochondrial bioenergetics and maintenance of mitochondrial membrane potential are important for energy production, limiting oxidative stress, and calcium buffering capacity. In these studies, we found mitochondrial bioenergetics was reduced acutely after SCI; however, ETC complex activities were not significantly different from control at 2h post-SCI. There were also no changes in mitochondrial dynamics or OXPHOS protein expression at this time point. This could indicate that mitochondrial respiratory dysfunction at 2h post-SCI was occurring before gross tissue damage was detected by ETC activities and Western blot. A previous report found mitochondrial bioenergetic dysfunction occurred at 6h post-SCI in female rats, which indicates that these present studies are the earliest account of mitochondrial impairment following SCI [17]. This information adds to our understanding of secondary injury following SCI, which can potentially be correlated to long-term outcomes and aid therapeutic development.

Contrary to the current literature, we did not find bioenergetic impairment 24h post-SCI in these studies. However, these studies were performed in male pigs, while the literature has primarily reported results in female rodents [17, 22, 29]. This could indicate there are differences in the time course of mitochondrial dysfunction between species or between sexes. Mitochondrial bioenergetics is represented as oxygen consumption rate (OCR), which could also indicate oxygen is being consumed by other factors, such as reactive oxygen species (ROS) generation. Interestingly, we did find an increase in ETC Complex Activities 24h post-SCI, which is consistent with increased protein expression of Complexes I and IV seen in these studies, as well. This, paired with decreased expression of the mitochondrial fusion protein, OPA1, suggests that there would be more fragmented mitochondria. However, fragmented mitochondria typically have a lower respiratory capacity and ETC complex activities, so the literature would suggest that other infiltrating cells, like neutrophils or microglia, contribute to increased ETC complex activity observed here [33, 34]. The OXPHOS antibody cocktail (Thermo Fisher) used here was designed to measure protein expressions of all 4 proteins of the ETC, as well as ATP Synthase, in one sample. The antibody for Complex V specifically targeted the alpha subunit of ATP Synthase, which is located on the F_1_ subunit. Interestingly, the F_1_ subunit has been shown to dissociate from F_0_ under pathological conditions, such as injury, which would result in increased proton leak; however, State IV respiration was not increased at this time point [35]. Contradicting data at this time point indicates that additional research is needed to understand the specific molecular sequelae and how this influences long-term recovery.

Most of what is known about mitochondrial dysfunction has been limited to acute and subacute time points following SCI [17, 21, 22, 29]. As SCI is a life-long condition, these studies performed mitochondrial analyses at 9wks to establish the mitochondrial profile at a chronic time point following SCI to improve long-term outcomes for those living with SCI. We found that State III respiration, also known as ATP production-linked respiration, was increased 9wks post-SCI, while no other respiration state was affected. Additionally, there were no changes in ETC complex activities, which is consistent with OXPHOS protein expression seen at this time point. There was also decreased Drp1 expression and increased PGC1α expression. Altogether, this suggests that mitochondrial dynamics have shifted toward recovery and energy production. This could be representative of the stability of astrocytes in actively maintaining the glial scar chronically following SCI [36, 37]. Oxidative stress markers associated SCI pathophysiology such as 3-NT, PC, and MDA, also demonstrated unique patterns of expression in pigs. The higher level of 3-NT, a marker of RNS damage, in SCI models in these studies relative to the uninjured controls is consistent with the prior studies in rats [38]. On the other hand, no differences in the expression of protein carbonyl, the ROS damage marker, across different time points in pig SCI models. However, PC is known to be increased in neurodegenerative conditions [39]. While MDA, the marker of lipid peroxidation, has been established to be higher in the plasma of human patients with chronic SCI than healthy control [40], it is robustly decreased in pigs with sub-acute (24h) SCI.

There are some limitations to mention about this study, with the first being small sample sizes. These studies were initially conducted in a preliminary subset of animals (n=3/group) for the 2h time point. We used State III data to estimate the sample size for each subsequent time point and calculated n=4 per group (G*Power; two-tailed unpaired t test, actual power = 0.96, effect size = 3.22). However, we were limited by a lack of both control and SCI animals for the 9wk bioenergetics experiments, so we maintained n=3/group for this preliminary study. Additionally, this study is limited by the time points selected for analysis. The 2h and 9wk time points were selected for the purposes of other studies, so mitochondrial analyses were added to the outcome measures to maximize data collection for each animal. The 24h time point was selected for this study because this time point is reported most often for mitochondrial analyses post-SCI and there is evidence of significant bioenergetic impairment and oxidative stress [16, 17, 21, 22]. However, mitochondrial dysfunction at 24h post-SCI has been primarily reported in female rodents, whereas, in these studies, we found no bioenergetic impairment at 24h post-SCI in male pigs. Additional research is needed to determine if this is due to differences in species or differences in sex. It is our intention to replicate this study in female pigs to investigate mitochondrial dysfunction across time following SCI to address the previous limitation.

This preliminary study was designed to establish a relative time course of mitochondrial function in an animal model with high translational fidelity. These experiments expand on the current literature which has primarily focused on acute time points, rodent models, and the female sex. By using male pigs and measuring mitochondrial function at a chronic time point after contusion/compression SCI, we aim to bridge the translational gap to understand secondary injury and improve the quality of life of those living with SCI.

## Acknowledgements

The authors would like to acknowledge Tracy Niedzielko and Jacob Jones for their assistance in surgery and tissue collection.

## References

1. Sezer, N., S. Akkuş, and F.G. Uğurlu, Chronic complications of spinal cord injury. World J Orthop, 2015. 6(1): p. 24–33.

2. Anderson, K.D., Targeting recovery: priorities of the spinal cord-injured population. J Neurotrauma, 2004. 21(10): p. 1371–83.

3. Higano, J.D., et al., Why do different people with Spinal Cord Injury have differing severity of symptoms with Autonomic Dysreflexia? Exploring relationships of vascular alpha-1 adrenoreceptor and baroreflex sensitivity after SCI. medRxiv, 2024: p. 2024.05.02.24306772.

4. Dhindsa, M.S., et al., Muscle spasticity associated with reduced whole-leg perfusion in persons with spinal cord injury. J Spinal Cord Med, 2011. 34(6): p. 594–9.

5. Khaing, Z.Z., et al., Clinical Trials Targeting Secondary Damage after Traumatic Spinal Cord Injury. Int J Mol Sci, 2023. 24(4).

6. Srikandarajah, N., M.A. Alvi, and M.G. Fehlings, Current insights into the management of spinal cord injury. J Orthop, 2023. 41: p. 8–13.

7. Hu, X., et al., Spinal cord injury: molecular mechanisms and therapeutic interventions. Signal Transduct Target Ther, 2023. 8(1): p. 245.

8. Hubbard, W.B., et al., Clinically relevant mitochondrial-targeted therapy improves chronic outcomes after traumatic brain injury. Brain, 2021. 144(12): p. 3788–3807.

9. Sullivan, P.G., et al., Mitochondrial uncoupling as a therapeutic target following neuronal injury. J Bioenerg Biomembr, 2004. 36(4): p. 353–6.

10. Velmurugan, G.V., et al., LRP1 Deficiency Promotes Mitostasis in Response to Oxidative Stress: Implications for Mitochondrial Targeting after Traumatic Brain Injury. Cells, 2023. 12(10).

11. LoPachin, R.M., et al., Experimental spinal cord injury: spatiotemporal characterization of elemental concentrations and water contents in axons and neuroglia. J Neurophysiol, 1999. 82(5): p. 2143–53.

12. Hachem, L.D., et al., Excitotoxic glutamate levels drive spinal cord ependymal stem cell proliferation and fate specification through CP-AMPAR signaling. Stem Cell Reports, 2023. 18(3): p. 672–687.

13. Pivovarova, N.B. and S.B. Andrews, Calcium-dependent mitochondrial function and dysfunction in neurons. Febs j, 2010. 277(18): p. 3622–36.

14. Sullivan, P.G., et al., Cytochrome c release and caspase activation after traumatic brain injury. Brain Res, 2002. 949(1-2): p. 88–96.

15. Sullivan, P.G., et al., Mitochondrial permeability transition in CNS trauma: cause or effect of neuronal cell deathJ Neurosci Res, 2005. 79(1-2): p. 231–9.

16. Patel, S.P., et al., N-acetylcysteine amide preserves mitochondrial bioenergetics and improves functional recovery following spinal trauma. Exp Neurol, 2014. 257: p. 95–105.

17. Sullivan, P.G., et al., Temporal characterization of mitochondrial bioenergetics after spinal cord injury. J Neurotrauma, 2007. 24(6): p. 991–9.

18. Hubbard, W.B., et al., Mitochondrial uncoupling prodrug improves tissue sparing, cognitive outcome, and mitochondrial bioenergetics after traumatic brain injury in male mice. J Neurosci Res, 2018. 96(10): p. 1677–1688.

19. Pandya, J.D., J.R. Pauly, and P.G. Sullivan, The optimal dosage and window of opportunity to maintain mitochondrial homeostasis following traumatic brain injury using the uncoupler FCCP. Exp Neurol, 2009. 218(2): p. 381–9.

20. Pandya, J.D., et al., N-acetylcysteine amide confers neuroprotection, improves bioenergetics and behavioral outcome following TBI. Exp Neurol, 2014. 257: p. 106–13.

21. Patel, S.P., et al., Pioglitazone treatment following spinal cord injury maintains acute mitochondrial integrity and increases chronic tissue sparing and functional recovery. Exp Neurol, 2017. 293: p. 74– 82.

22. Patel, S.P., et al., Differential effects of the mitochondrial uncoupling agent, 2,4-dinitrophenol, or the nitroxide antioxidant, Tempol, on synaptic or nonsynaptic mitochondria after spinal cord injury. J Neurosci Res, 2009. 87(1): p. 130–140.

23. Gollihue, J.L., et al., Effects of Mitochondrial Transplantation on Bioenergetics, Cellular Incorporation, and Functional Recovery after Spinal Cord Injury. J Neurotrauma, 2018. 35(15): p. 1800–1818.

24. Gollihue, J.L., S.P. Patel, and A.G. Rabchevsky, Mitochondrial transplantation strategies as potential therapeutics for central nervous system trauma. Neural Regen Res, 2018. 13(2): p. 194–197.

25. Lin, M.W., et al., Mitochondrial Transplantation Attenuates Neural Damage and Improves Locomotor Function After Traumatic Spinal Cord Injury in Rats. Front Neurosci, 2022. 16: p. 800883.

26. Kalimon, O.J., et al., Characterizing Sex Differences in Mitochondrial Dysfunction After Severe Traumatic Brain Injury in Mice. Neurotrauma Rep, 2023. 4(1): p. 627–642.

27. Hill, R.L., et al., Time courses of post-injury mitochondrial oxidative damage and respiratory dysfunction and neuronal cytoskeletal degradation in a rat model of focal traumatic brain injury. Neurochem Int, 2017. 111: p. 45–56.

28. Singh, I.N., et al., Time course of post-traumatic mitochondrial oxidative damage and dysfunction in a mouse model of focal traumatic brain injury: implications for neuroprotective therapy. J Cereb Blood Flow Metab, 2006. 26(11): p. 1407–18.

29. Stewart, A.N., et al., Mitochondria exert age-divergent effects on recovery from spinal cord injury. Exp Neurol, 2021. 337: p. 113597.

30. Vekaria, H.J., et al., An efficient and high-throughput method for the evaluation of mitochondrial dysfunction in frozen brain samples after traumatic brain injury. Front Mol Biosci, 2024. 11: p. 1378536.

31. Musyaju, S., et al., Revert total protein normalization method offers a reliable loading control for mitochondrial samples following TBI. Anal Biochem, 2023. 680: p. 115301.

32. Yan, L.J. and M.J. Forster, Chemical probes for analysis of carbonylated proteins: a review. J Chromatogr B Analyt Technol Biomed Life Sci, 2011. 879(17-18): p. 1308–15.

33. Sterner, R.C. and R.M. Sterner, Immune response following traumatic spinal cord injury: Pathophysiology and therapies. Front Immunol, 2022. 13: p. 1084101.

34. Picard, M., et al., Mitochondrial morphology transitions and functions: implications for retrograde signalingAm J Physiol Regul Integr Comp Physiol, 2013. 304(6): p. R393–406.

35. Mnatsakanyan, N., et al., Mitochondrial ATP synthase c-subunit leak channel triggers cell death upon loss of its F(1) subcomplex. Cell Death Differ, 2022. 29(9): p. 1874–1887.

36. Gaudet, A.D. and L.K. Fonken, Glial Cells Shape Pathology and Repair After Spinal Cord Injury. Neurotherapeutics, 2018. 15(3): p. 554–577.

37. Tamaru, T., et al., Glial scar survives until the chronic phase by recruiting scar-forming astrocytes after spinal cord injury. Exp Neurol, 2023. 359: p. 114264.

38. Khan, M., et al., Amelioration of spinal cord injury in rats by blocking peroxynitrite/calpain activity. BMC Neurosci, 2018. 19(1): p. 50.

39. Shen, L., et al., Redox Proteomic Profiling of Specifically Carbonylated Proteins in the Serum of Triple Transgenic Alzheimer’s Disease Mice. Int J Mol Sci, 2016. 17(4): p. 469.

40. Haro Girón, S., et al., Prognostic Value of Malondialdehyde (MDA) in the Temporal Progression of Chronic Spinal Cord Injury. J Pers Med, 2023. 13(4).

